# Signatures of selective sweeps in continuous-space populations

**DOI:** 10.1101/2024.07.26.605365

**Authors:** Meera Chotai, Xinzhu Wei, Philipp W. Messer

## Abstract

Selective sweeps describe the process by which an adaptive mutation arises and rapidly fixes in the population, thereby removing genetic variation in its genomic vicinity. The expected signatures of selective sweeps are relatively well understood in panmictic population models, yet natural populations often extend across larger geographic ranges where individuals are more likely to mate with those born nearby. To investigate how such spatial population structure can affect sweep dynamics and signatures, we simulated selective sweeps in populations inhabiting a two-dimensional continuous landscape. The maximum dispersal distance of offspring from their parents can be varied in our simulations from an essentially panmictic population to scenarios with increasingly limited dispersal. We find that in low-dispersal populations, adaptive mutations spread more slowly than in panmictic ones, while recombination becomes less effective at breaking up genetic linkage around the sweep locus. Together, these factors result in a trough of reduced genetic diversity around the sweep locus that looks very similar across dispersal rates. We also find that the site frequency spectrum around hard sweeps in low-dispersal populations becomes enriched for intermediate-frequency variants, making these sweeps appear softer than they are. Furthermore, haplotype heterozygosity at the sweep locus tends to be elevated in low-dispersal scenarios as compared to panmixia, contrary to what we observe in neutral scenarios without sweeps. The haplotype patterns generated by these hard sweeps in low-dispersal populations can resemble soft sweeps from standing genetic variation that arose from substantially older alleles. Our results highlight the need for better accounting for spatial population structure when making inferences about selective sweeps.

## Introduction

The classic selective sweep model introduced by Smith and Haigh provides the theoretical foundation for many studies of adaptation in natural and experimental populations (Smith and Haigh (1974)). This model describes how positive selection can drive a *de novo* adaptive mutation to high frequency in a population and makes several predictions about the expected patterns of surrounding genetic diversity: As the adaptive mutation spreads, linked neutral variants can “hitchhike” with it, while non-adaptive haplotypes will be displaced from the population, thereby depleting genetic diversity (Smith and Haigh (1974)). In the absence of recombination, mutations occurring on the sweeping haplotype can create new variants of this haplotype, which should result in a characteristic power-law decay in the haplotype frequency spectrum when the adaptive mutation becomes fixed (Messer and Neher (2012)).

On a recombining chromosome, crossover events during the sweep can also generate new adaptive haplotype variants that preserve ancestral genetic variation. This should produce a characteristic trough in neutral genetic diversity around the sweep locus, with the site frequency spectrum (SFS) of neutral polymorphisms in this region displaying an excess of both high- and low-frequency derived alleles (Fay and Wu (2000); Kim and Stephan (2002)). The expected size of this trough is approximately proportional to the ratio of *s*/*r*, where *s* is the selection coefficient of the adaptive mutation and *r* is the recombination rate along the chromosome (Kaplan *et al*. (1989)). Linkage disequilibrium (LD) and haplotype homozygosity should both be elevated around the sweep locus as compared to a neutrally evolving region in equilibrium (Hudson *et al*. (1994)). These predicted sweep signatures lie at the core of many existing methods for the detection and study of selective sweeps from population genomic data (Messer and Neher (2012); DeGiorgio *et al*. (2016); Schrider and Kern (2016); Hejase *et al*. (2021)).

A key assumption underlying the classic sweep model is that populations are homogeneously mixed with randomly mating individuals (so-called “panmixia”). However, natural populations are often structured in various ways, which can lead to dramatic violations of the panmixia assumption (Durrett and Levin (1994)). Many real-world populations inhabit a continuous geographic range, for example, where individuals tend to disperse over distances much shorter than the dimensions of the habitat. Individuals will thus be more likely to mate with individuals who were born nearby. This can increase levels of inbreeding and produce noticeably different patterns of neutral genetic diversity as compared to a panmictic population (Wright (1943); Barton *et al*. (2002); Ringbauer *et al*. (2017); Battey *et al*. (2020); Etheridge *et al*. (2023)).

Spatial structure can also significantly affect the spread of an adaptive mutation when the dispersal of individuals becomes a limiting factor (Barton *et al*. (2013)). In such limited-dispersal scenarios, a strongly advantageous mutation cannot rise in frequency as quickly as it would in a panmictic population, and its frequency trajectory will deviate from the logistic growth curve expected under the panmictic sweep model (Maynard Smith (1971)). Instead, these dynamics have commonly been described by Fisher traveling waves (Fisher (1937)). In a one-dimensional (1D) space, the wave model predicts that the frequency of the adaptive mutation should increase approximately linearly over time like the distance traveled by a wave that moves outward from its origin at constant velocity; in 2D space, it should increase approximately quadratically like the area of a circle with a linearly growing radius. These disparities in sweep dynamics could impact various sweep signatures, such as the establishment probability of the adaptive mutation, the fixation time of the sweep, and the expected patterns in surrounding genetic diversity.

This raises a key question: How accurate is the classic selective sweep model for studying adaptation in real-world populations that inhabit continuous space? Min *et al*. (2022) recently explored this question for 1D space using the abstraction of a stepping-stone model, wherein the population is subdivided into discrete panmictic demes along a one-dimensional array with migration occurring only between neighboring demes (Kimura and Weiss (1964)). They found that sweep signatures under such a model differ from the classic sweeps in several ways. Most notably, the SFS of neutral variants around the sweep locus should exhibit a flat tail with many intermediate frequency variants consisting of mutations that fixed in the wavefront of the sweep and then hitchhiked through the rest of the population (Min *et al*. (2022)). This is very different from the classic sweep model where the SFS around the sweep locus should be heavily skewed towards both low- and high-frequency variants (Fay and Wu (2000)).

Interestingly, an excess of intermediate frequency variants in the SFS is commonly associated with so-called “soft” selective sweeps, which occur when not all adaptive lineages have coalesced at the onset of positive selection (Hermisson and Pennings (2005); Prezeworski *et al*. (2005); Messer and Petrov (2013); Hermisson and Pennings (2017)). Such soft sweeps can occur when adaptive alleles were either already present as standing genetic variation (SGV) at the onset of selection, or when adaptive alleles arose recurrently during the sweep. By contrast, sweeps involving only a single *de novo* adaptive mutation, as assumed by the classic selective sweep model, are commonly referred to as “hard” selective sweeps.

Whether hard or soft selective sweeps are more common in nature has been intensely debated over the past years (Karasov *et al*. (2010); Jensen (2014); Feder *et al*. (2016); Hermisson and Pennings (2017); Schrider and Kern (2017); Garud *et al*. (2021)). To tackle this question, various summary statistics and computational inference methods have been developed for discriminating among the different types of sweeps (Prezeworski *et al*. (2005); Peter *et al*. (2012); Ferrer-Admetlla *et al*. (2014); Garud *et al*. (2015); Schrider and Kern (2016)). However, these approaches typically assume a panmictic population. It is therefore critical to know whether patterns of diversity left behind by hard sweeps in continuous-space populations can indeed resemble those of soft sweeps in panmictic models, and thereby perhaps confound our inferences. While Min *et al*. (2022) focused on SFS-based signatures, it remains unclear whether the apparent “softening” of sweep signatures in continuous-space populations extends to patterns beyond the SFS that can discriminate hard from soft selective sweeps more effectively, such as haplotype structure (Ferrer-Admetlla *et al*. (2014); Garud *et al*. (2015)). Furthermore, we do not know how results from a 1D model translate to 2D continuous space, which should be more appropriate for the majority of natural populations.

In this paper, we conduct a comprehensive analysis of selective sweeps in continuous-space populations. Using individual-based simulations of a population that inhabits a 2D continuous-space landscape, we study how different dispersal rates affect sweep trajectories, establishment probabilities, and fixation times for varying selection strengths, as compared to a panmictic population model with otherwise similar properties. We then examine how these changes in sweep dynamics – as well as local versus global sampling schemes – impact the observed patterns of neutral genetic variation around the sweep locus, including the size of the trough in neutral diversity, the SFS, and the haplotype frequency spectrum.

## Model

We implemented an individual-based simulation model in SLiM version 4.0.1 (Haller and Messer (2023)) to study selective sweeps in 2D spatial populations of diploid, sexually reproducing hermaphrodites. Our model is designed to allow for a gradual transition from a panmictic population to one with strongly limited dispersal, while otherwise resembling a standard Wright-Fisher model as closely as possible. In particular, we assume discrete, non-overlapping generations, and a constant population size (Walsh and Lynch (2018)). We also want to keep the local population density constant over time and space. This is a non-trivial setup for a spatial model, as fluctuations in local density can often arise due to the random nature of dispersal, mating, and other stochastic processes (Felsenstein (1975); Sasaki (1997); Vinatier *et al*. (2011)). In addition, the requirement of a constant local density entails a model of “soft” selection (Wallace (1975)), in which selection only affects local allele frequencies but not the overall number of individuals in any given region of the population (as compared to any model of “hard” selection, which directly affects viability and/or fecundity).

Our motivation for seeking a soft selection model with constant population density is grounded in several factors: First, such a model should more closely resemble standard theoretical results, most notably the Fisher-KPP equation for modeling the spatial dynamics of an adaptive allele in a reaction-diffusion framework (Fisher (1937)). Second, a hard selection model would typically require some form of local density control to avoid runaway clustering (Felsenstein (1975)), introducing a level of complexity we would like to avoid here. Finally, the definition of a selection coefficient can be more ambiguous in a hard selection model, rendering key properties such as the establishment probability or fixation time of a sweep more difficult to compare with theoretical expectations.

To satisfy all of the above assumptions, we implemented our simulation model as follows: Individuals inhabit a continuous-space 2D square arena of size 1 × 1. The arena features periodic, “toroidal” boundaries, where the edges connect seamlessly to the opposite edges (e.g., an individual that exits the upper-left corner returns in the lower-right one). This allows our model to avoid various edge effects encountered in scenarios with reflective, absorbing, or reprising boundaries (Mazzucco *et al*. (2018)). Generations are discrete and non-overlapping. In each generation, a new set of *N* = 10^4^ individuals is generated with parents drawn from the individuals in the previous generation. To achieve locally uniform population density, we imposed a 100 × 100 square grid on our square arena, such that exactly one individual always lives inside each grid cell (Figure 1A). The coordinates of each individual are chosen randomly within its cell, and its two parents are randomly sampled (without replacement) from those individuals in the previous generation that were located within a circle of radius *d*, centered at the location of the newly generated individual. The dispersal radius *d* thereby specifies the maximum possible distance between offspring and parent. In the following, we will always refer to *d* as the “dispersal rate” of the model. Once all offspring are generated, the individuals from the previous generation are removed and the generation cycle starts anew.

**Figure 1.**
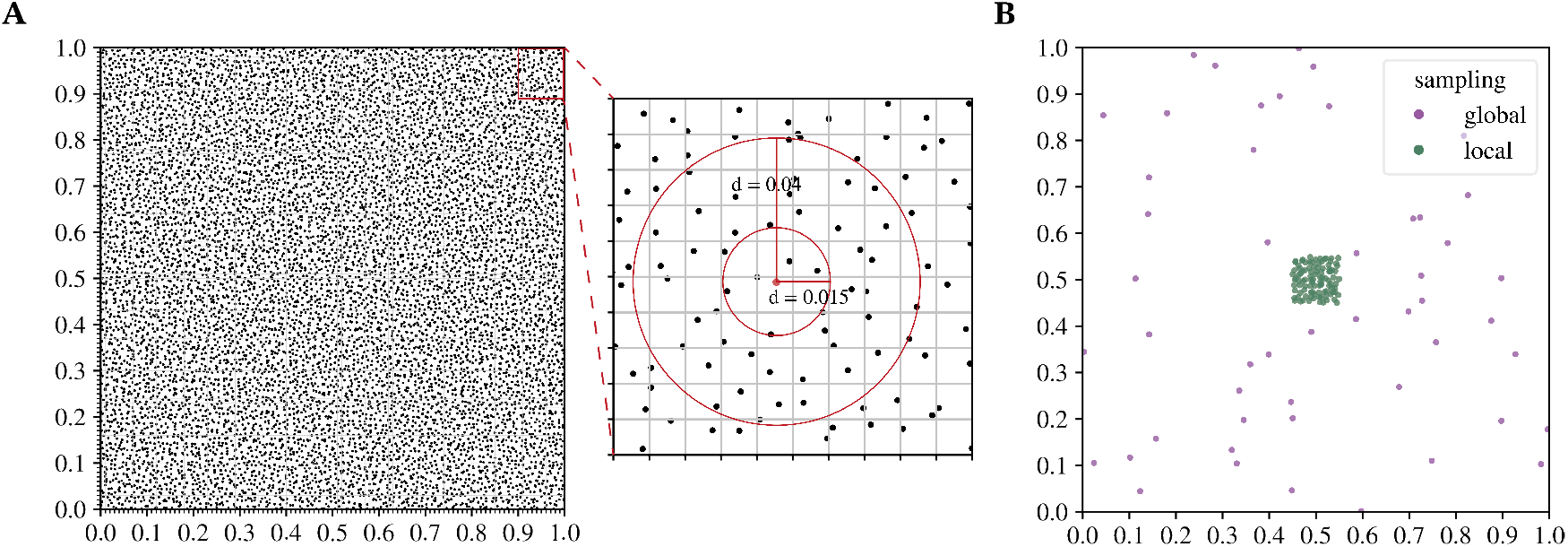
Simulation model and sampling strategy. A. We simulate a 2D continuous-space square arena of size 1 × 1 with toroidal boundaries, inhabited by *N* = 104 individuals (black dots). To ensure constant population size and uniform population density, the arena is split into a 100 × 100 square grid, with one individual living per grid cell in each generation. The zoom into the upper right corner shows how a focal individual (red dot) picks its two parents from the individuals of the previous generation (black dots). Two dispersal rates *d* = 0.015, 0.04 are shown (red circles), each containing all potential parents for the focal individual under the given value of *d*. **B**. Illustration of the local and global sampling strategies used for the evaluation of sweep signatures. A global sample (shown in purple) contains individuals randomly drawn from the entire arena, while a local sample (shown in green) contains only individuals randomly drawn from within a square region of size 0.1 × 0.1 in the center of the arena.

By varying the dispersal rate *d*, we can gradually transition from a scenario with highly localized dispersal to a completely panmictic population. We chose a value of *d* = 0.015 as the low-dispersal limit, which ensures that for any newly generated individual, two parents can practically always be found within its dispersal radius. By contrast, the high-dispersal limit was set at *d* = 1.0, where the circle encompasses all individuals of the previous generation, thus corresponding to a completely panmictic population.

Under this model, we simulated the evolution of a chromosome of length 10 Mbp with a recombination rate of 10^−8^ per bp per generation and a neutral mutation rate of 10^−8^ mutations per bp per generation, with neutral mutations being overlaid onto tree sequences obtained at the end of each simulation run using msprime (version 1.2.0) (Baumdicker *et al*. (2021)).

For each value of *d* used in our simulations, we first performed three replicate neutral burn-in simulations runs of 5 × 10^5^ generations to allow the simulated population to reach equilibrium (confirmed by testing that 100% of sites have successfully coalesced at the end of the simulation run). The resulting populations were then output as tree sequence files (Kelleher *et al*. (2018); Haller *et al*. (2019)).

Although our spatial model does not encompass all aspects of natural populations in continuous space, it captures key aspects of spatiality by allowing for dispersal and variation in local allele frequencies across the habitat, while at the same time controlling for other potentially complicating factors such as fluctuations in total population size and local density. We therefore believe that our model can serve as a useful baseline model for understanding the effects of continuous space and dispersal on the dynamics and signatures of selective sweeps.

### Sweep simulations

To simulate a selective sweep, we first load a randomly chosen neutral burn-in tree sequence file for the given dispersal rate *d* into SLiM and then introduce an adaptive allele of the given selection coefficient *s* at the center of a randomly sampled chromosome from the population. We assume codominance (*h* = 1/2), such that wildtype homozygotes have fitness 1.0, heterozygotes have fitness 1 + *s*/2, and homozygotes for the adaptive mutation have fitness 1 + *s*.

Selection is implemented in our model by weighing the probabilities of picking different parents within the dispersal circle of a newly generated individual according to their relative fitness values. For example, if three individuals are present inside the circle, with two of them having fitness 1.0 while the third has fitness 1.2 (because it is homozygous for an adaptive mutation with selection coefficient *s* = 0.2), the first two individuals would each be picked with probability 1.0/3.2 while the third would be picked with probability 1.2/3.2. Since we sample without replacement, if we end up picking one of the individuals with fitness 1.0 as the first parent, the probabilities for picking the second parent would then be 1.0/2.2 and 1.2/2.2, respectively. In the panmictic limit, where the dispersal circle encompasses all individuals in the population, this approach should converge to a standard Wright-Fisher model with selection.

Each selective sweep simulation was run until either fixation or loss of the adaptive mutation. If the mutation reached fixation, we stored the tree sequence file and then overlaid neutral mutations onto the tree sequences to obtain the resulting patterns of neutral diversity (Haller *et al*. (2019); Kelleher *et al*. (2018); Ralph *et al*. (2020b)).

For the softness analysis presented in Figure 7, we simulated soft sweeps from SGV in a panmictic Wright-Fisher model, where the adaptive allele was previously neutral and present at a given starting frequency (*x*_0_) in the population before it became adaptive. These sweeps were modeled by first simulating a neutral equilibrium population with Θ = 4*N*_*e*_*µ* = 0.0004 in the coalescent simulator msprime (version 1.2.0) (Baumdicker et al. 2021). We then randomly chose one of the neutral mutations that had the required derived allele frequency and was located at least 50 kbp away from the ends of the genome. This mutation was subsequently assigned a selection coefficient of *s* = 0.1, while all other neutral mutations were removed and the simulation was saved as a tree-sequence file. The tree-sequence files were then loaded into SLiM and a forward-in-time simulation was performed until eventual fixation or loss of the now-beneficial allele. The coalescent and forward-in-time simulations were repeated until we had 1000 fixation events for each desired starting frequency, which were all saved as tree-sequence files. Finally, neutral mutations were overlaid onto the tree sequences to obtain the resulting patterns of neutral diversity, analogous to the hard sweep simulations.

### Local versus global sampling

Generating a population sample can be more complicated in a spatial population as compared to a panmictic one because the location of individuals could be a factor in the sampling strategy. We studied two distinct sampling strategies in our spatial model: global versus local sampling (illustrated in Figure 1B). To generate a global sample, 50 individuals (and thus, *n* = 100 chromosomes) are randomly drawn from the entire population (without replacement). By contrast, a local sample is generated by drawing 50 individuals from within a square region of side-length 0.1 in the center of the arena. The individuals in a local sample therefore lived much closer to each other as compared to those from a global sample. Note that in our sweep simulations, we chose the origin of each sweep randomly across the entire habitat. Thus, the central habitat area from which we draw local samples usually does not coincide with the area in which the sweep originated, so this sampling strategy can still be considered random relative to the sweep origin.

### Estimation of summary statistics

For Figure 2, we computed several summary statistics on populations after the neutral burn-in. For each dispersal rate *d*, we used one replicate to obtain a distribution of coalescence times estimated across all sites of the simulated chromosome for 50 pairs of individuals in a global sample. Variation in coalescence time is directly reflected in the measurement of nucleotide heterozygosity (*π*), defined as the average number of heterozygous sites in a pair of chromosomes normalized by its length (Nei and Li (1979)). We used tskit to measure the average pairwise neutral heterozygosity (*π*_0_) for local and global samples of individuals, showing the mean over all three replicates (Kelleher *et al*. (2018); Ralph *et al*. (2020a,b)). In addition, we calculated the average neutral heterozygosity within diploid genomes for a sample of individuals, showing the mean over all three replicates.

**Figure 2.**
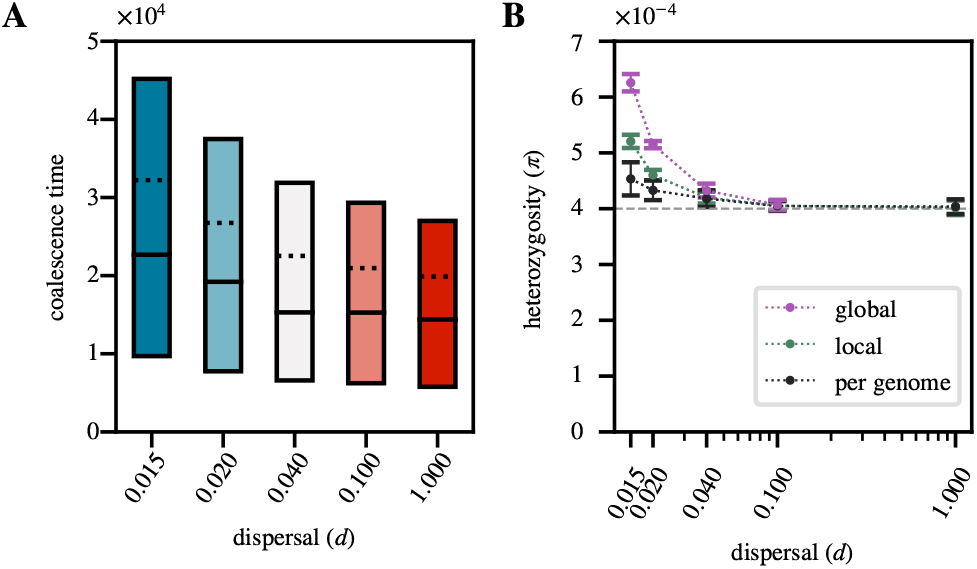
Patterns of neutral diversity under limited dispersal: A. Distribution of pairwise coalescence times in neutral simulations for different dispersal rates after a burn-in of 50*N* generations. The solid lines show the 25th, 50th and 75th percentiles, while the dotted line shows the mean. **B**. Nucleotide heterozygosity in neutral simulations for different dispersal rates in global samples, local samples, and when estimated in a diploid genome. The grey dashed line represents the panmictic expectation of *π* ≈ 4*Nµ*. The mean and standard deviations (represented by the error bars) were calculated from 100 random samples across the 3 neutral replicates.

**Figure 3.**
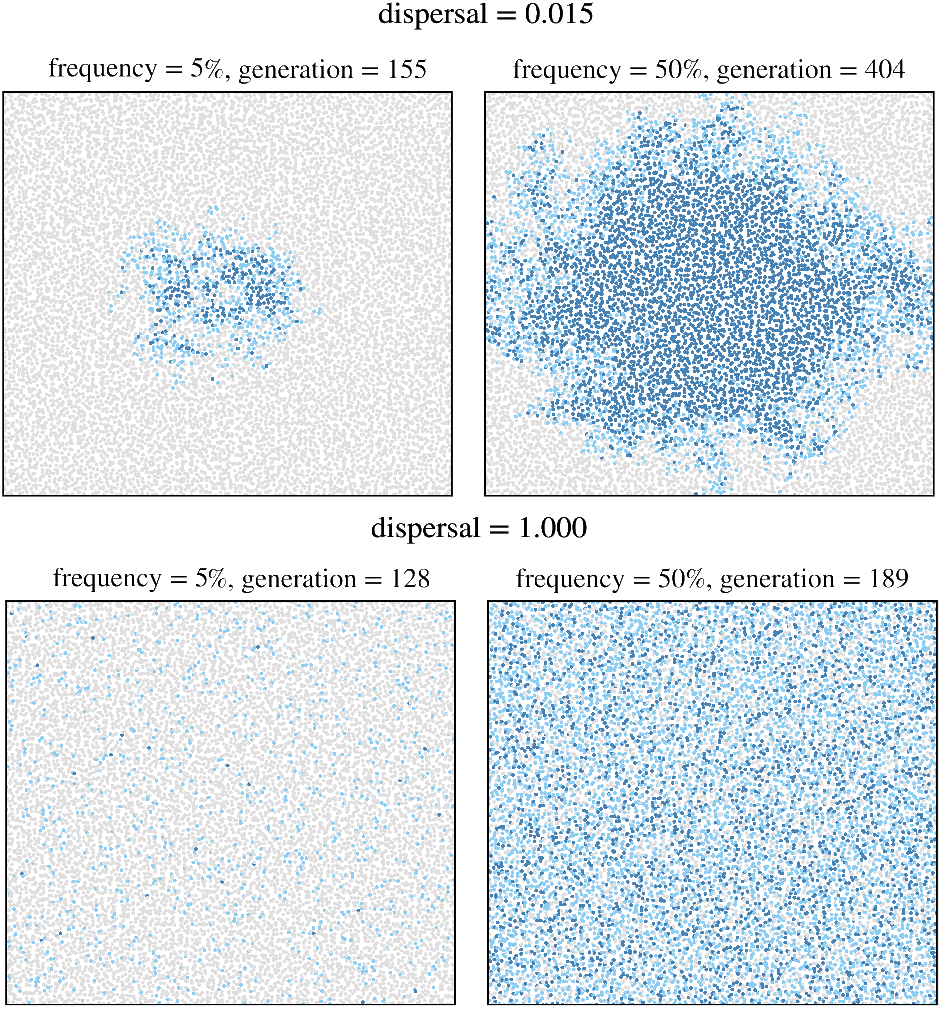
Qualitative illustration of sweep dynamics under limited-dispersal (top row, *d* = 0.015) versus panmixia (bottom row, *d* = 1.0). Each row shows two population snapshots, taken when the adaptive mutation first reached a population frequency of 5% (left) and 50% (right). Light blue points represent heterozygous individuals, while dark blue points represent homozygous individuals. The adaptive mutation had a selection coefficient of *s* = 0.1 and was introduced close to the center of the habitat.

**Figure 4.**
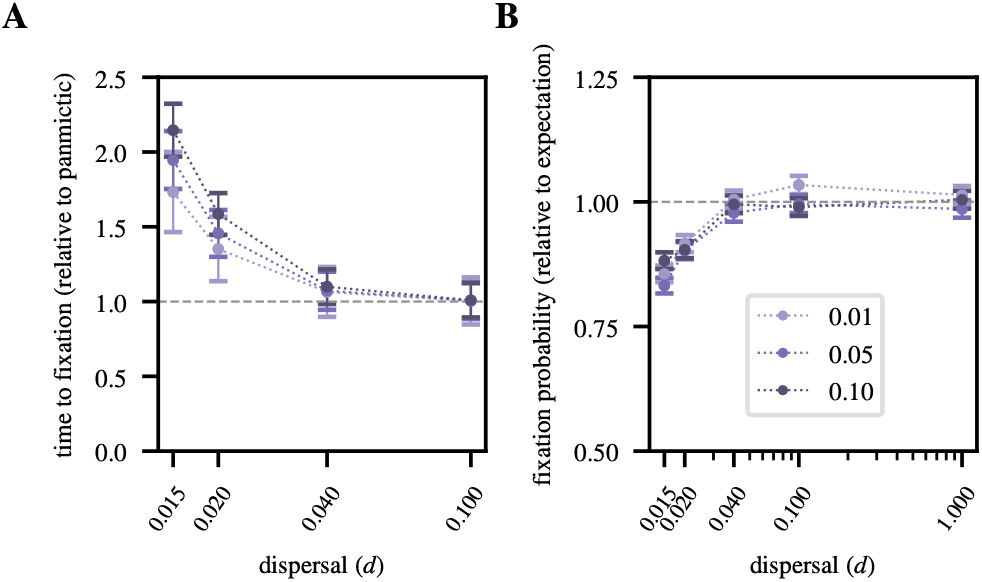
Fixation times and probabilities: A. Ratio of the average sweep fixation time for varying dispersal rates relative to the average fixation time under panmixia. Error bars represent the standard deviation from the mean, estimated over ∼3000 successful sweeps per data point. **B**. Ratio of the fixation probability of a new adaptive mutation with selection coefficient *s* for varying dispersal rates relative to the expected fixation probability under panmixia ((1 −*e*−*s*)/(1 −*e*−2*Ns*)). Each data point was estimated from ∼ 3000 successful sweeps. Error bars represent the binomial sampling error, calculated as 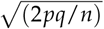

For Figures 5A and 6A-B, we computed several summary statistics using individuals sampled from the population immediately after the sweeps reached fixation. We used tskit to compute the heterozygosity, SFS, and Tajima’s *D* within a genomic window centered at the sweep locus (Kelleher *et al*. (2018); Ralph *et al*. (2020a,b)). The sizes of the windows were chosen to capture the characteristic trough in heterozygosity around the sweep locus, and therefore depend on the strength of selection.

Specifically, we chose a window size of 150 kbp for *s* = 0.01 and 1 Mbp for *s* = 0.1. For all three sweep statistics, we obtained the mean and standard deviation over 1000 replicate simulations for each combination of *s* and *d*. For comparison, we also computed the SFS and Tajima’s *D* across the entire chromosome for individuals from the neutral burn-in simulations, averaging over all three replicates. In addition, for Figure 5B we identified runs of homozygosity (ROHs), using PLINK (version 1.9) with command-line arguments “–homozyg-snp 20 –homozyg-kb 200 –homozyg-het 0” (Purcell and Chang (2022); Chang *et al*. (2015)). We calculated the average proportion of a diploid genome covered by ROHs for global samples of individuals, averaging over the three neutral burn-in replicates.

For Figures 6C-D, we calculated several haplotype statistics for individuals sampled from populations immediately after the sweeps reached fixation. For Figure 6C, we calculated the average haplotype frequency spectra for global and local samples of individuals for the low-dispersal limit (*d* = 0.015) and global samples of individuals for the panmictic limit (*d* = 1.0), using 1000 replicates each and considering sweeps of strength *s* = 0.1. We recorded the frequency of distinct haplotypes, and arranged them in descending order of frequency (Miles *et al*. (2023)). For Figures 6D and E, we derived the haplotype heterozygosity and the number of haplotypes across the range of dispersal rates and selection strengths of *s* = 0.01, 0.1. Haplotype heterozygosity is defined as the probability that two randomly sampled chromosomes are distinct, calculated as 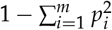, where *m* is the total number of distinct haplotypes at the time of fixation in a given sample set, and *p*_*i*_ is the frequency of the haplotype *i* in the sample set (Muralidhar and Veller (2022)). These statistics were computed over a range of genomic window sizes (10 kbp, 30 kbp, and 100 kbp) centered around the sweep locus, and were averaged over 1000 replicates for each data point. We calculated the haplotype heterozygosity and the number of haplotypes for samples from the neutral burn-in simulations, arbitrarily centering the windows around the center of the genome and averaging over all three replicates. Additionally, for Figure 7 we calculated the haplotype heterozygosity for the soft SGV sweeps within a window of size 100 kbp (the largest of our three window sizes) centered at the sweep locus. We computed the average haplotype heterozygosity over 1000 replicates for each value of *x*_0_.

**Figure 5.**
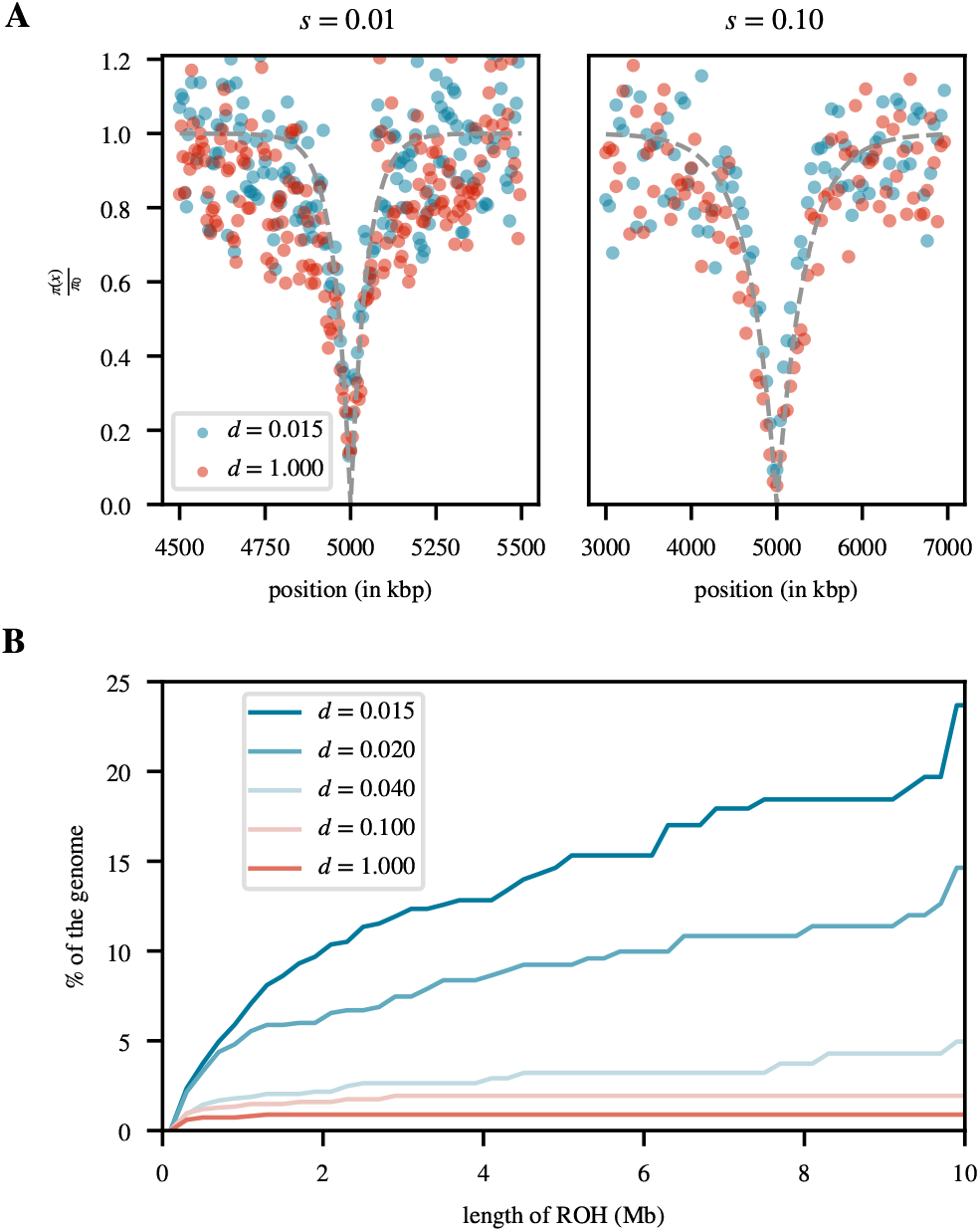
Sweep sizes and ROH distributions: A. Reduction in heterozygosity (*π*) around the sweep locus estimated at sweep fixation relative to the neutral level before the sweep (*π*_0_) for the two extreme dispersal rates *d* = 0.015 and *d* = 1.0, and for selection strengths of *s* = 0.01 (left) and *s* = 0.1 (right). For the left plot, each data point was estimated over a non-overlapping window of size 5 kbp, averaged over 1000 replicates. For the right plot, each data point was estimated over a non-overlapping window of size 40 kbp, averaged over 1000 replicates. The grey dashed line represents the panmictic expectation of 1− (2*Ns*)^−4*rx*/*s*^, where *x* is the distance from the sweep locus (Barton (2000)). **B**. Cumulative proportion of a diploid genome covered by ROHs as a function of the maximum length of ROHs considered. For each value of *d*, the result represents an average across samples from three neutral burn-in replicates. A minimum length of 200 kbp is used for calling ROHs.

**Figure 6.**
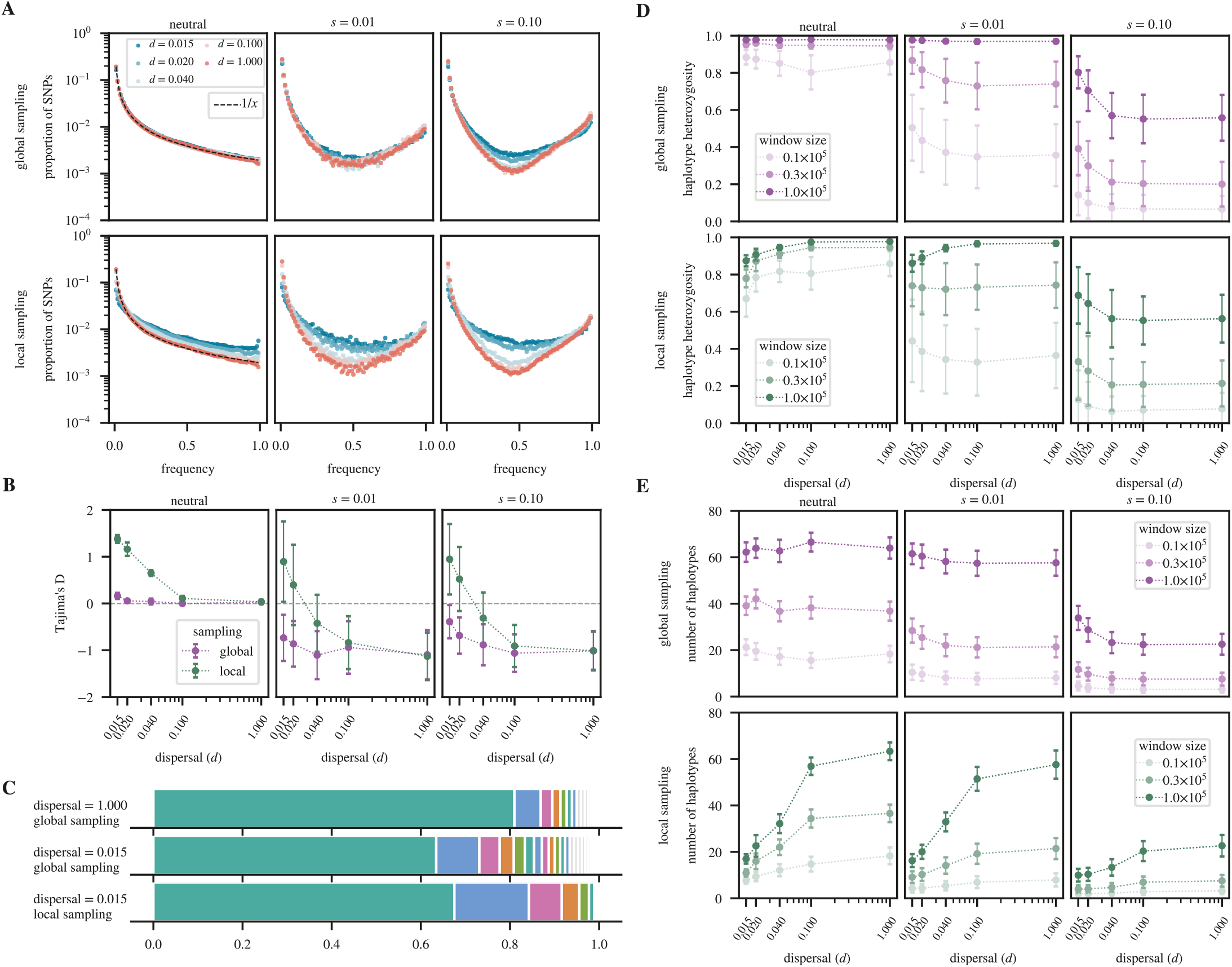
Softening of hard selection sweeps under limited dispersal: A. Site frequency spectra under neutrality and two sweep strengths for varying dispersal rates. For the neutral simulations, the spectra were estimated across the entire simulated chromosome of length 10 Mbp. For the sweep scenarios, the windows were centered on the sweep locus, with their sizes chosen to capture the average dip in diversity (150 kbp for *s* = 0.01, and 1 Mbp for *s* = 0.1). **B**. Tajima’s *D* values for each of the spectra estimated above. Values of *D* = 0 correspond to the neutral equilibrium expectation (grey dashed line); negative values indicate a skew towards low- and/or high-frequency derived variants; positive values indicate enrichment for intermediate-frequency variants. **C**. Haplotype frequency spectra for selective sweep simulations of strength *s* = 0.1 within a window of size 50 kbp centered on the sweep for the panmictic limit (*d* = 1.0) and the low-dispersal limit (*d* = 0.015), and estimated in global versus local samples. Each haplotype frequency spectrum was averaged over 1000 sweeps. **D**. Haplotype heterozygosity under neutrality and two selection strengths for varying dispersal rates, different window sizes, and global versus local sampling. **E**. Same as **D**, but showing the over-all number of haplotypes present in the analysis window instead of haplotype heterozygosity. The mean and standard deviation (represented by the error bars) of each data point in **B**,**D**,**E** were estimated across 1000 replicates.

**Figure 7.**
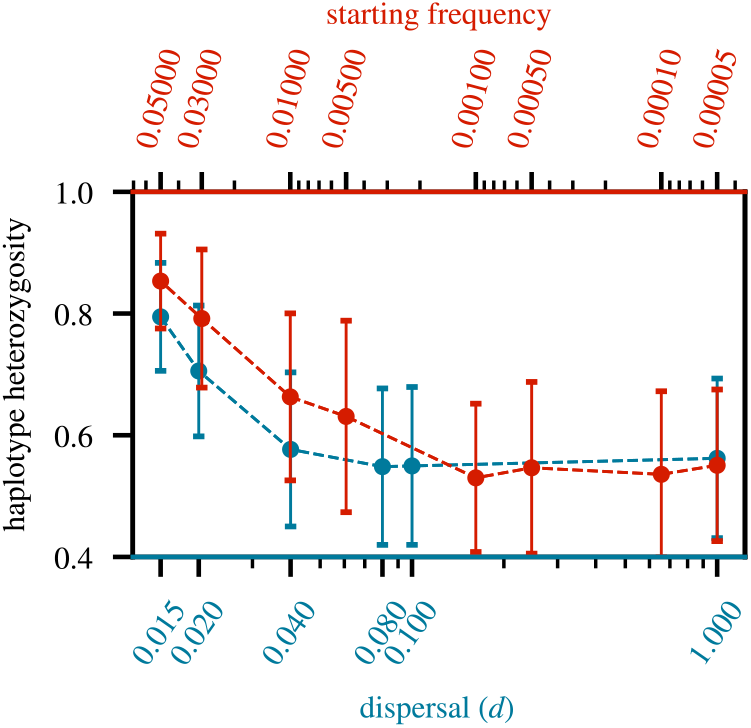
Softness comparison between SGV and spatial sweeps: The two curves show average haplotype heterozygosity levels measured in windows of size 100 kbp in global samples for sweeps of strength *s* = 0.1. The blue curve depicts the results for hard sweeps in our spatial model for varying dispersal rates (bottom x-axis), while the red curve depicts the results for SGV sweeps in a panmictic population for varying starting frequencies (top x-axis). Note that the top x-axis is inverted. Higher haplotype heterozygosity indicates a softer sweep, as found for SGV sweeps with higher starting frequencies and spatial hard sweeps in scenarios with lower dispersal rates. The mean and standard deviation (represented by the error bars) of each data point were estimated across 1000 replicates.

## Results

Our individual-based simulation model allows us to study evolutionary dynamics in continuous-space populations with varying levels of dispersal, ranging from an effectively panmictic population to one where offspring are displaced from their parents by an average distance of just 1-2% of the diameter of the simulated habitat. The parameter choices for our baseline model (*N* = 10^4^, *µ* = *r* = 10^−8^) are broadly inspired by a regime relevant to human populations, with an expected pairwise coalescence time of 2*N* = 20, 000 generations and an equilibrium nucleotide heterozygosity level of *π* ≈ 4*Nµ* = 0.0004 when assuming panmixia. Under this model, we simulated the evolution of a chromosomal region of length 10 Mbp.

First, we wanted to assess the impact of spatiality on equilibrium patterns of neutral genetic diversity for varying dispersal rates *d*, as this pre-existing diversity provides the background on which sweeps will act. The results shown in Figure 2 confirm that limited dispersal increases pairwise coalescence times and thus also *π*. This known effect is due to the characteristic changes in genealogical histories in spatial populations with limited dispersal, where spatially distant genomes will have deeper coalescence histories as compared to two randomly sampled genomes from an equally sized panmictic population (Charlesworth *et al*. (2003); Ralph and Coop (2013)). The elevation is more pronounced in global as compared to local samples, since the latter will more frequently contain highly related individuals with shorter branches among them. The smallest increase is observed when *π* is estimated in one diploid genome, which can be considered a “maximally local” sample of two haploid chromosomes. In the panmictic limit, pairwise coalescence time and nucleotide heterozygosity in our simulation converge to their theoretical panmictic expectations regardless of the sampling strategy.

### Sweep dynamics in the spatial population model

Next, we studied how limited dispersal in a spatial population affects the dynamics of hard selective sweeps from a single *de novo* adaptive mutation. Figure 3 shows examples of two simulated sweep runs originating at the center of the arena that already illustrate three qualitative differences in the sweep dynamics between a spatial population with limited dispersal and a panmictic one: (i) in the spatial population with limited dispersal, the sweep spreads in a wave-like manner from its point of origin, whereas under panmixia, the frequency of the adaptive mutation increases homogeneously across the habitat; (ii) in the spatial population with limited dispersal, the adaptive allele spreads slower as compared to panmixia; (iii) in the spatial population with limited dispersal, heterozygotes are present only at the wavefront, and their relative frequency in the population tends to be lower throughout the sweep as compared to panmixia. These effects are generally consistent with previous results (Barton *et al*. (2013); Min *et al*. (2022)). Below, we will investigate how they quantitatively impact sweep dynamics and the resulting signatures in neutral genetic diversity.

The reason for the prolonged fixation times in a spatial model with limited dispersal can be understood when considering the expected change in the population frequency *x*(*t*) of the adaptive mutation. For a co-dominant mutation (*h* = 1/2) with selection coefficient *s* in a panmictic Wright-Fisher population, these dynamics are described by the logistic differential equation *dx*(*t*)/*dt* ≈ *sx*(1− *x*)/2, where the approximation is accurate as long as *sx* ≪ 1 (Kimura (1962)). The rate at which the adaptive mutation can spread is thus limited solely by its selection coefficient. In a continuous-space population with limited dispersal, by contrast, the maximum rate of spread can also be limited by the dispersal of individuals. Regardless of selection strength, the adaptive mutation can only spread as far in each generation as offspring are dispersed from their parents. When *d* becomes small enough, this can become the limiting factor for the rate of spread, resulting in a prolonged fixation time.

Figure 4A shows the magnitude of this effect in our spatial model for different values of *d*. For instance, adaptive mutations with *s* = 0.01 have an approximately 1.7 × longer average fixation time in the low-dispersal limit (*d* = 0.015) compared to their average fixation time under panmixia. This effect is more pronounced the larger the selection coefficient (e.g., for sweeps with *s* = 0.1 the average fixation time is approximately 2.1× longer in the low-dispersal limit compared to the panmictic limit).

A slower rate of spread of an adaptive mutation under limited dispersal could have important consequences during the early sweep phase. In particular, if a mutation remains at a low frequency for a longer time, there should be an increased probability of stochastic loss due to genetic drift, decreasing the overall fixation probability of such mutations. In a panmictic diploid population of size *N*, a new mutation with selection coefficient *s* and dominance coefficient *h* = 1/2 initially present in one copy is expected to ultimately become fixed in the population with probability (1 − *e*^−*s*^)/(1− *e*^−2*Ns*^) (Kimura (1957)). Our simulations confirm this classic theoretical result in the panmictic limit (Figure 4B). For limited dispersal scenarios, fixation probabilities indeed slightly decrease from the panmictic expectations. For instance, the probability of fixation of a new mutation with *s* = 0.1 in the low-dispersal limit is ∼ 20% lower than under panmixia.

### Limited dispersal has little effect on sweep size

A key signature of a selective sweep is the reduction of genetic diversity in the vicinity of the sweep locus due to hitchhiking. At fixation, the expected size of this trough is determined by the probability that recombination events occurring during the sweep were able to break up linkage between the adaptive alleles and linked neutral hitchhiker alleles. A higher recombination rate and/or a longer fixation time should therefore produce a narrower trough, and vice versa. Under panmixia, expected fixation times are roughly proportional to the selection coefficient of the sweep, so sweep sizes should be approximately proportional to *s*/*r* (Kaplan *et al*. (1989)). Several existing methods leverage this result to estimate the selection strengths of recently completed sweeps (Kim and Stephan (2002); Sattath *et al*. (2011)).

Previous studies have suggested that the increased fixation time in a spatial population with limited dispersal should lead to narrower sweep sizes compared to panmixia (Kim and Maruki (2011); Barton *et al*. (2013); Min *et al*. (2022)). Thus, methods for estimating selection strength based on sweep size could underestimate the true *s*. We investigated whether this prediction holds true in our spatial population model. Contrary to expectation, we found that sweep sizes are in fact very similar in the low-dispersal and panmictic limits, with no discernible differences across the range of selection coefficients we tested (Figure 5A).

One hypothesis for why this could be the case is that recombination might be less effective at breaking up genetic linkage in low-dispersal populations, thereby compensating for the longer fixation times. In particular, we surmised that the presence of long ROHs, which should be more common in populations with low dispersal due to higher inbreeding rates (Min *et al*. (2022)), could be lowering the efficacy of recombination. This is because any crossover event occurring inside an ROH will be ineffective at shuffling alleles at other loci within the same ROH. In other words, looking backward in time, if the two segments created by a recombination breakpoint coalesce with each other before either coalesces with another lineage, the recombination event has no effect on linkage patterns within that region (Nordborg (2000)).

Indeed, we observed a dramatic increase in the number of long ROHs for populations under limited dispersal as compared to panmixia (Figure 5B). For example, in panmictic populations, ROHs (defined as homozygous regions of length ≥ 200 kbp containing ≥ 20 SNPs, see Model) make up only ∼ 0.89% of the cumulative genome length of a sample of *n* = 50 individuals, while this number is approximately 26 times higher (∼ 23.7%) in the low-dispersal limit (*d* = 0.015). In comparison to other summary statistics of neutral diversity, such as *π*, this is arguably the most striking difference we observed between the panmictic and low-dispersal limits. However, while this substantial increase in long ROHs should lower the efficacy of recombination, it is not clear whether this effect is truly a driving factor in determining sweep size, and why it would almost perfectly counteract the effect of longer fixation times. A more rigorous study of this question would require a better understanding of how continuous spatial population structure with limited dispersal affects linkage patterns and the effectiveness of recombination in breaking up linkage when long ROHs are common.

### Limited dispersal can make sweeps appear softer

In the hard selective sweep model implemented in our simulations, a sweep arises from a single *de novo* mutation that is immediately adaptive. Under panmixia, this model predicts the classic sweep signatures: a reduction in *π* around the sweep locus, an excess of high- and low-frequency derived nucleotide variants in the SFS, and the presence of one long haplotype at high population frequency. By contrast, soft selective sweeps from SGV or recurrent *de novo* mutations tend to produce markedly different signatures. In particular, nucleotide heterozygosity around the sweep locus is not necessarily reduced to the same extent as in a hard sweep, the SFS can show an excess of intermediate-frequency variants, and several long haplotypes can be present at intermediate population frequencies, resulting in generally higher levels of haplotype heterozygosity at the sweep locus (Hermisson and Pennings (2017)).

Based on the data depicted in Figure 5A, the reduction in *π* around the sweep locus looks very similar in the panmictic and low-dispersal limits. At the center of the sweep, *π* diminishes to nearly 0% of its neutral equilibrium value in either scenario, suggesting that limited dispersal does not make these hard sweeps appear softer in that regard. This is consistent with the fact that at the sweep locus, all lineages present when the adaptive mutation fixes must have coalesced by the time this mutation first arose. Even though it may take the adaptive mutation about twice as long to go to fixation in the low-dispersal limit as compared to pamixia, this is not enough to substantially increase levels of neutral diversity close to the sweep locus.

A different result has been suggested for SFS-based signatures. In particular, Min *et al*. (2022) modeled hard selective sweeps in a 1D stepping-stone model and found an excess of intermediate frequency variants in the SFS compared to the classic sweep model, which led them to conclude that hard selective sweeps in 1D spatially structured populations may appear softer than they actually are. To investigate whether we can also observe this “softening” effect in our 2D continuous-space model, we studied the SFS and Tajima’s *D* values estimated around simulated sweeps in global samples (Figures 6A-B). In the panmictic limit, our simulations show the predicted excess of high- and low-frequency variants resulting in negative Tajima’s D values. However, as the dispersal rate decreases, the SFS indeed becomes enriched for intermediate-frequency SNPs, confirming that the apparent softening effect also occurs in our 2D continuous-space model. We further find that this effect is much more pronounced for stronger (*s* = 0.1) than weaker (*s* = 0.01) sweeps. In neutral simulations, SFS and Tajima’s *D* values closely follow the equilibrium expectation under panmixia across the range of dispersal rates when estimated in global samples.

In addition to nucleotide diversity and the SFS, haplotype-based signatures can provide a third distinguishing feature between hard and soft sweeps. In a hard sweep, when the adaptive mutation becomes fixed, the haplotype on which it originally arose should be the most prevalent one, while variants of this haplotype that arose from mutation or recombination events during the sweep should typically remain at much lower frequencies. By contrast, soft sweeps can involve several different adaptive haplotypes present at intermediate frequencies, resulting in generally higher levels of haplotype heterozygosity around the sweep locus.

Indeed, we find that haplotype frequency spectra around hard sweeps (with *s* = 0.1) also appear softer under limited dispersal (Figure 6). When estimating the average haplotype frequency spectrum in a window of size 50 kbp around such sweeps in global samples, we observe a clear shift towards more intermediate frequency haplotypes in the low-dispersal limit as compared to panmixia. This is consistent with a systematic increase in haplotype heterozygosity around the sweep locus under limited dispersal (Figure 6D), while no such increase is observed under neutrality. Finally, there is also an increase in the overall number of haplotypes in the analysis window, especially for the stronger sweeps (Figure 6E). Again, no such increase is observed under neutrality.

### Effect of local sampling

Next, we wanted to examine how these findings are affected when we estimate SFS and haplotype statistics within local samples rather than global ones. In this case, the enrichment of intermediate frequency variants in the SFS around sweeps due to limited dispersal is even more pronounced (Figure 6A). For example, while Tajima’s D values estimated in global samples remain negative for almost all simulated sweeps across the range of dispersal regimes tested, in local samples they are almost always positive in the low-dispersal limit, with typical values on the order of one (Figure 6B). We also observe an enrichment of intermediate frequency variants under limited dispersal in neutral simulations in local samples, while no such effect was seen in global samples. This is consistent with previous studies that observed a shift in the SFS towards intermediate frequencies in local samples taken from neutrally evolving, spatially-structured populations (De and Durrett (2007); Städler *et al*. (2009); Min *et al*. (2022)).

Similar to what we observed in global samples, haplotype heterozygosity generally increases in local samples with lower dispersal rates, except for the weak sweeps (*s* = 0.01) in the longest windows (100 kbp), where we see a slight decrease in haplotype heterozygosity in the low-dispersal limit. This decrease is also observed in neutral simulations, presumably because of longer and more frequent ROHs in such samples. Thus, for weak sweeps, our longest windows presumably contain enough neutral diversity far away from the sweep to revert to the patterns observed under neutrality. As demonstrated in Figure 6E, there is a marked decrease in the overall number of different haplotypes observed within local samples, thus revealing no clear signature of softening. However, this effect is also observed under neutrality.

Overall, these findings suggest that limited dispersal can produce a softening effect in haplotype signatures around the sweep locus, marked by increased haplotype heterozygosity (in both global and local samples) and higher haplotype numbers (in global but not local samples). In most cases, these effects do not conform to the impact of limited dispersal on haplotype patterns in neutral scenarios.

### Comparing spatial hard sweeps to SGV soft sweeps

One scenario that can produce true soft sweeps is when adaptation occurs from SGV. For instance, a previously neutral allele could have already been present at a certain starting frequency in the population when it suddenly became adaptive. To better understand the degree by which limited dispersal softens the hard sweeps in our spatial model, we wanted to investigate which starting frequency in this panmictic SGV soft sweep model yields similar levels of haplotype heterozygosity to hard sweeps in our spatial model under limited dispersal. When the starting frequency is very low, the level of haplotype heterozygosity observed at fixation of SGV soft sweeps should converge to that of spatial hard sweeps in the panmictic limit, while systematically increasing with higher starting frequency.

Figure 7 shows the haplotype heterozygosity of simulated soft sweeps from SGV in a panmictic population as a function of their starting frequency, compared to the haplotype heterozygosity of simulated hard sweeps in spatial populations as a function of the dispersal rate. The right end of the plot depicts the panmictic limit of the spatial model, and SGV sweeps with the lowest possible starting frequency *x*_0_ = 1/(2*N*), corresponding to a single copy of the adaptive allele. In this limit, both curves converge to a haplotype heterozygosity of ∼ 0.56, as expected from Figure 6D. On the left end of the plot, which depicts the low-dispersal limit of the spatial hard sweep model and SGV sweeps with high starting frequency, we find that haplotype heterozygosity in the spatial hard sweeps is elevated to ∼ 0.794. This is comparable to the haplotype heterozygosity of 0.791 measured for SGV soft sweeps with a starting frequency of *x*_0_ = 0.03. Thus, for this specific scenario, limited dispersal softens haplotype heterozygosity levels of hard sweeps in our spatial model to an extent where they resemble soft sweeps from SGV in a panmictic population starting from a previously neutral allele that was initially present in 2*Nx*_0_ = 600 copies when it became adaptive.

One conceivable explanation for this observed degree of softening could be the increased fixation time in a low-dispersal scenario as compared to panmixia. In other words, maybe the panmictic SGV sweeps that yield comparable levels of haplotype heterozygosity as the spatial hard sweeps under limited dispersal have adaptive alleles with comparable overall ages at fixation as these hard sweeps, and this essentially explains why they appear softer. However, as we already showed above, a mutation with *s* = 0.1 undergoing a hard sweep in a low-dispersal scenario only takes about twice as long to fix (∼ 730 generations) as a mutation undergoing a hard sweep in a panmictic population (∼ 340 generations). By contrast, the mean age of a neutral mutation present at a starting frequency of *x*_0_ = 0.03 is ∼ 3500 generations in our SGV sweep simulations, which is close to the theoretical expectation (Kimura and Ohta (1973)). The total average age of such a mutation at the fixation of an SGV soft sweep is then ∼ 3800 generations, and thus they are substantially older than their counterparts in the spatial hard sweep model, suggesting that the increase in fixation time alone cannot explain the elevated haplotype heterozygosity.

## Discussion

Our current understanding of selective sweeps is largely based on panmictic models where all individuals in the population are well mixed. This assumption could be problematic for many real-world organisms that inhabit larger geographic ranges but only disperse over short distances during their lifetime. In this study, we used individual-based simulations of selective sweeps to investigate how such spatial structure can affect sweep dynamics and signatures in 2D continuous-space populations.

We first showed that limited dispersal can slow down the spread of an adaptive allele and thereby lower its fixation probability compared to expectations in a panmictic population of equal size, echoing previous findings in a 1D stepping-stone model (Min *et al*. (2022)). In the low-dispersal regime, the population dynamics of the adaptive allele are no longer described by logistic growth, but rather by a Fisher traveling wave spreading in an approximately circular fashion outward from the point of origin of the adaptive allele. As a result, heterozygotes are mostly found at the wavefront of the expanding sweep, and their population frequency deviates substantially from the Hardy-Weinberg equilibrium. We showed that these qualitatively different dynamics can profoundly alter the signatures left by a selective sweep in surrounding genetic diversity from those expected under panmixia. In many cases, the effects reflect a complex interplay between the changed sweep dynamics and the impact of limited dispersal on equilibrium patterns of neutral genetic variation that provide the background on which the sweep occurs.

For example, we showed that contrary to what had been predicted in previous studies (Min *et al*. (2022); Barton *et al*. (2013)), the slowed spread of the sweep under low dispersal does not necessarily result in a narrower trough of neutral diversity around the sweep locus. This is surprising because a longer fixation time should allow for more recombination events during the sweep that could break up the linkage between the adaptive allele and surrounding neutral variants. It is, therefore, not entirely clear why we do not observe a narrower trough in our simulations. We surmised that this may be due to a reduced effectiveness of recombination from crossover events occurring inside ROHs, which are much longer and more prevalent in low-dispersal populations as compared to panmictic ones, presumably due to higher rates of local inbreeding. A crossover event within an ROH cannot shuffle alleles at other loci within the same ROH (Nordborg (2000)). Thus, if ROHs are common and long enough to extend over typical sweep sizes, this should lower the effective rate of recombination between SNPs within those distances, while not affecting the rates across loci that are farther apart.

The consequences of a reduction in the effective recombination rate on selection have been studied extensively in the context of selfing populations Hedrick (1980); Charlesworth *et al*. (1997); Hartfield and Glémin (2014); Roze (2016); Hartfield and Bataillon (2020); Sianta *et al*. (2022). Indeed, several of these studies showed that a reduced effective recombination rate in selfing populations is expected to increase the span of the hitch-hiking region in a selective sweep. More rigorous methods are still needed to understand how these results could apply to continuous-space populations. Nevertheless, if our finding that sweep size is not severely affected by low dispersal does hold more generally, this would imply that methods for inferring selection strength based on this signal remain robust in low-dispersal populations even when their inferences rely on predictions from panmictic models.

Regarding the SFS around a sweep locus, our study revealed that low-dispersal causes an enrichment of intermediate frequency SNPs compared to the panmictic expectation. A similar phenomenon was noted by Min *et al*. (2022) for selective sweeps in a 1D stepping-stone model. They proposed that this flattening of the SFS results from mutations that fix in the wavefront of the sweep and are then carried through the rest of the population, while also suggesting that this effect might be less pronounced in 2D populations. Our results demonstrate that the flattening effect is still clearly observable in 2D populations with limited dispersal.

Furthermore, we found that low dispersal can systematically alter the haplotype structure observed around a sweep locus, with characteristic changes in the haplotype frequency spectrum. Specifically, we showed that limited dispersal generally increases the level of haplotype heterozygosity around the sweep locus, contrary to what is observed under neutrality. This is true for local and global samples but with different underlying changes in the haplotype frequency spectrum in each case. In global samples, the average number of haplotypes observed in a genomic window increases under low dispersal, both around a sweep locus and under neutrality (consistent with the longer coalescence times in low-dispersal populations). By contrast, average haplotype numbers decrease in local samples in both scenarios (consistent with the higher prevalence of long ROHs). Yet we still observed higher haplotype heterozygosity around a sweep with lower dispersal even in local samples because more of those haplotypes tend to be present at intermediate frequencies.

A key question is whether the parameter regime we studied could be relevant to real-world populations so that these effects can manifest in practice. Generally, we expect those effects to become more severe with stronger selection and lower dispersal rate (relative to the extent of the population’s geographic range). In our simulation model, we studied sweeps with selection coefficients ranging from *s* = 0.01 to *s* = 0.1. These are arguably relatively strong sweeps. However, many examples of sweeps of such strength have been observed in nature. For instance, the selection coefficient of the sweep associated with lactase persistence in humans has been estimated to be in the range of 1 − 2 percent (Bersaglieri *et al*. (2004)). Even stronger sweeps have been observed at the MC1R gene in Monarch birds associated with color pigmentation (Campagna *et al*. (2022)), or the evolution of pesticide resistance in *Anopheles* mosquitoes (Lynd *et al*. (2010)). There is evidence that sweeps with selection coefficients on the order of 1% or larger could have been common in various species (Sella *et al*. (2009); Enard *et al*. (2014); Rogers *et al*. (2023); Wei *et al*. (2023)). Thus, we conclude that the strength of selection modeled in our study is not unrealistic.

Neither should the dispersal rates studied in our model be considered unrealistically low for many real-world populations. We observed the strongest effects in our model’s low-dispersal limit, which was set at *d* = 0.015. This means that the average distance between parent and offspring locations is still roughly one percent of the diameter of the population range. In many real-world populations, dispersal rates should actually be lower. For example, the average distance between the birthplaces of second-degree cousins or closer was found to be only < 5 km in the UK Biobank (Nait Saada *et al*. (2020)), which is significantly less than one percent of the diameter of the UK. Similarly, the maximum recorded dispersal distance in *Anopheles gambiae* mosquitoes was found to be less than 700 meters in a mark–release–recapture study in Kenya (Midega *et al*. (2007)), again a much shorter distance than one percent of this species’ geographic range in sub-Saharan Africa.

Hence, we believe that the effects of limited dispersal on sweep dynamics and signatures we demonstrated in this study could be relevant for various real-world populations inhabiting large geographical ranges, such as humans or mosquitoes. One indication that this might be the case for any given sweep would be the observation of clear differences in haplotype patterns around the sweep locus between local and global samples, as illustrated in Figure 6E.

An important caveat to our study is that we did not include the potential for long-range dispersal, which could be an important factor in shaping the dynamics of sweeps in real-world populations (Paulose *et al*. (2019)). When the dispersal kernel has a long tail such that a few individuals occasionally migrate over much larger distances than typical, this could lead to multiple introductions of the adaptive allele into distant regions of the population. Each such new introduction can potentially seed a new patch in which the allele can then spread locally, accelerating its overall spread (Paulose *et al*. (2019)).

Our model makes several other simplifications that could limit its generality. In particular, we modeled a completely homogeneous population with a constant local density and dispersal rate across space and time. Real-world populations will always be more heterogeneous, and this could affect sweep behavior in various ways. We also only studied a population of a fixed size of *N* = 10, 000 individuals, neglecting any potential impact of demographic events such as historic bottlenecks or expansions. A larger or smaller population size (while keeping the dispersal rate and habitat size the same) will change the rate of genetic drift in the population. This could impact both the background patterns of neutral diversity and the dynamics of the adaptive allele in complex ways that remain to be explored.

Importantly, we also assumed a highly idealized sweep model involving a codominant adaptive mutation with a constant selection coefficient. These assumptions are likely violated at least to some extent in most real-world populations. For recessive mutations in particular, sweep dynamics could be even more different in the continuous-space model as compared to a panmictic population because when the mutation is still rare, homozygotes can be present in the population at a higher frequency than expected under Hardy-Weinberg proportions, thereby potentially facilitating its initial spread.

Perhaps most problematic is our assumption of a population sample obtained precisely when the adaptive mutation fixes in the population. In most practical applications, the population will likely be sampled when the sweep is either still ongoing or has already ended. In fact, many sweeps may not reach fixation at all due to various factors, such as when the benefit of the adaptive allele weakens over time or exists only in certain contexts or parts of the population (Pritchard *et al*. (2010)). The signatures produced by such partial selective sweeps can be more multi-faceted than the simple model we studied here, as both adaptive and non-adaptive haplotypes will be captured in the sample.

Given all these simplifications, as well as the general difficulty of parameterizing any real-world spatial population model, our results should not be seen as questioning the results from any particular sweep inference method. However, they do raise the possibility that spatial population structure could be a strong confounding factor in some previous studies that assumed panmixia, especially when selection is strong. In particular, the flattening of the SFS we observed in low-dispersal populations, in combination with the increase in haplotype heterozygosity, are both commonly associated with soft selective sweeps. Some methods for distinguishing hard from soft sweeps based on such signals could hence be misled into classifying hard sweeps as soft if they rely on expectations from panmictic models.

Our results highlight a general problem for methods that base their inferences solely on summary statistics, which can often yield the same results under fundamentally different evolutionary scenarios. For sweep classification, the defining feature is the true genealogy of adaptive alleles at the selected site. In a hard sweep, by definition, all adaptive lineages in the sample must coalesce more recently than when it first became advantageous to carry the allele. A soft sweep, by contrast, is defined by coalescence occurring before the onset of positive selection (Messer and Petrov (2013); Hermisson and Pennings (2017)). All sweeps we simulated in our spatial model were hard by definition, arising from a single mutation that was immediately adaptive. If the true genealogy at the selected site were known, it would be clear that these sweeps were hard, and did not arise from a mutation in the SGV that had been segregating for thousands of generations before it became adaptive, as suggested by the observed level of haplotype heterozygosity. We hope that recent progress in the development of methods for accurate inference of ancestral recombination graphs (Brandt *et al*. (2024); Deng *et al*. (2024)) could pave the way for a new class of sweep inference methods that are more robust to confounding factors than current approaches that rely on summary statistics alone.

The appeal of the classic selective sweep model introduced by Smith and Haigh stems in no small part from its elegance and simplicity, yet biological reality is inevitably more complex. Consequently, there has been a long history of studying how various potentially confounding factors, such as background selection and demography, can affect selective sweeps and potentially bias the inferences of methods that do not account for them appropriately. Our study adds another layer of complexity to this challenge by demonstrating that continuous spatial population structure can also profoundly impact sweep dynamics and signatures.

In light of our findings, we suggest that methods for studying selective sweeps, when applied to populations where dispersal could be a limiting factor, should explicitly consider the potential impact that such spatial structure might have on their inferences.

In some cases, it may be evident that the level of dispersal in the study population is high enough over the timescales relevant to a sweep that the assumption of a panmictic model remains justified. In other cases, it may be necessary to explicitly show that results obtained under this assumption remain robust for a given level of population structure. When this is not the case, spatial structure should be explicitly included in the model or analysis. This can be challenging because of the complexity involved in accurately parameterizing spatial models, where key parameters such as the dispersal kernel of the study population are often unknown. However, with the advent of powerful simulation tools (Chevy *et al*. (2024)), it is now at least feasible to simulate evolutionary dynamics in baseline models of continuous-space populations that can help us better understand how such structure affects evolutionary processes, as we have demonstrated in this study.

## Data availability

Scripts used for all analyses and figures are available at https://github.com/meerachotai/spatial_selection.

## Acknowledgments

The authors thank Ben Haller, Sam Champer, and members of the Wei and Messer labs for helpful discussions.

## Literature cited

Barton N, Etheridge A, Kelleher J, Véber A. 2013. Genetic hitch-hiking in spatially extended populations. Theoretical Population Biology. 87:75–89.

Barton NH. 2000. Genetic hitchhiking. Philosophical Transactions of the Royal Society of London. Series B: Biological Sciences. 355:1553–1562.

Barton NH, Depaulis F, Etheridge AM. 2002. Neutral Evolution in Spatially Continuous Populations. Theoretical Population Biology. 61:31–48.

Battey CJ, Ralph PL, Kern AD. 2020. Space is the Place: Effects of Continuous Spatial Structure on Analysis of Population Genetic Data. Genetics. 215:193–214.

Baumdicker F, Bisschop G, Goldstein D, Gower G, Ragsdale AP, Tsambos G, Zhu S, Eldon B, Ellerman EC, Galloway JG et al. 2021. Efficient ancestry and mutation simulation with msprime 1.0. Genetics. 220:iyab229.

Bersaglieri T, Sabeti PC, Patterson N, Vanderploeg T, Schaffner SF, Drake JA, Rhodes M, Reich DE, Hirschhorn JN. 2004. Genetic signatures of strong recent positive selection at the lactase gene. The American Journal of Human Genetics. 74:1111–1120.

Brandt DY, Huber CD, Chiang CW, Ortega-Del Vecchyo D. 2024. The Promise of Inferring the Past Using the Ancestral Recombination Graph. Genome Biology and Evolution. 16:evae005.

Campagna L, Mo Z, Siepel A, Uy JAC. 2022. Selective sweeps on different pigmentation genes mediate convergent evolution of island melanism in two incipient bird species. PLOS Genetics. 18:1–25.

Chang CC, Chow CC, Tellier LC, Vattikuti S, Purcell SM, Lee JJ. 2015. Second-generation PLINK: rising to the challenge of larger and richer datasets. GigaScience. 4:s13742.–015–0047–8.

Charlesworth B, Charlesworth D, Barton NH. 2003. The Effects of Genetic and Geographic Structure on Neutral Variation. Annual Review of Ecology, Evolution, and Systematics. 34:99–125.

Charlesworth B, Nordborg M, Charlesworth D. 1997. The effects of local selection, balanced polymorphism and background selection on equilibrium patterns of genetic diversity in subdivided populations. Genetics Research. 70:155–174.

Chevy ET, Min J, Caudill V, Champer SE, Haller BC, Rehmann CT, Smith CCR, Tittes S, Messer PW, Kern AD et al. 2024. Population genetics meets ecology: a guide to individual-based simulations in continuous landscapes. bioRxiv.

De A, Durrett R. 2007. Stepping-Stone Spatial Structure Causes Slow Decay of Linkage Disequilibrium and Shifts the Site Frequency Spectrum. Genetics. 176:969–981.

DeGiorgio M, Huber CD, Hubisz MJ, Hellmann I, Nielsen R. 2016. SweepFinder2: increased sensitivity, robustness and flexibility. Bioinformatics. 32:1895–1897.

Deng Y, Nielsen R, Song YS. 2024. Robust and Accurate Bayesian Inference of Genome-Wide Genealogies for Large Samples. bioRxiv.

Durrett R, Levin S. 1994. The Importance of Being Discrete (and Spatial). Theoretical Population Biology. 46:363–394.

Enard D, Messer PW, Petrov DA. 2014. Genome-wide signals of positive selection in human evolution. Genome Research. 24:885–895.

Etheridge AM, Kurtz TG, Letter I, Ralph PL, Lung TTH. 2023. Looking forwards and backwards: dynamics and genealogies of locally regulated populations.

Fay JC, Wu CI. 2000. Hitchhiking Under Positive Darwinian Selection. Genetics. 155:1405–1413.

Feder AF, Rhee SY, Holmes SP, Shafer RW, Petrov DA, Pennings PS. 2016. More effective drugs lead to harder selective sweeps in the evolution of drug resistance in HIV-1. eLife. 5:e10670.

Felsenstein J. 1975. A Pain in the Torus: Some Difficulties with Models of Isolation by Distance. The American Naturalist. 109:359–368.

Ferrer-Admetlla A, Liang M, Korneliussen T, Nielsen R. 2014. On Detecting Incomplete Soft or Hard Selective Sweeps Using Haplotype Structure. Molecular Biology and Evolution. 31:1275–1291.

Fisher RA. 1937. The Wave of Advance of Advantageous Genes. Annals of Eugenics. 7:355–369.

Garud NR, Messer PW, Buzbas EO, Petrov DA. 2015. Recent Selective Sweeps in North American Drosophila melanogaster Show Signatures of Soft Sweeps. PLOS Genetics. 11:1–32.

Garud NR, Messer PW, Petrov DA. 2021. Detection of hard and soft selective sweeps from drosophila melanogaster population genomic data. PLOS Genetics. 17:1–35.

Haller BC, Galloway J, Kelleher J, Messer PW, Ralph PL. 2019. Tree-sequence recording in SLiM opens new horizons for forward-time simulation of whole genomes. Molecular Ecology Resources. 19:552–566.

Haller BC, Messer PW. 2023. SLiM 4: Multispecies EcoEvolutionary Modeling. The American Naturalist. 201:E127–E139.

Hartfield M, Bataillon T. 2020. Selective Sweeps Under Dominance and Inbreeding. G3 Genes|Genomes|Genetics. 10:1063–1075.

Hartfield M, Glémin S. 2014. Hitchhiking of Deleterious Alleles and the Cost of Adaptation in Partially Selfing Species. Genetics. 196:281–293.

Hedrick PW. 1980. Hitchhiking: A Comparison of Linkage and Partial Selfing. Genetics. 94:791–808.

Hejase HA, Mo Z, Campagna L, Siepel A. 2021. A Deep-Learning Approach for Inference of Selective Sweeps from the Ancestral Recombination Graph. Molecular Biology and Evolution. 39:msab332.

Hermisson J, Pennings PS. 2005. Soft Sweeps: Molecular Population Genetics of Adaptation From Standing Genetic Variation. Genetics. 169:2335–2352.

Hermisson J, Pennings PS. 2017. Soft sweeps and beyond: understanding the patterns and probabilities of selection footprints under rapid adaptation. Methods in Ecology and Evolution. 8:700–716.

Hudson RR, Bailey K, Skarecky D, Kwiatowski J, Ayala FJ. 1994. Evidence for positive selection in the superoxide dismutase (Sod) region of Drosophila melanogaster. Genetics. 136:1329– 1340.

Jensen JD. 2014. On the unfounded enthusiasm for soft selective sweeps. Nature Communications. 5:1–10.

Kaplan NL, Hudson RR, Langley CH. 1989. The “hitchhiking effect” revisited. Genetics. 123:887–899.

Karasov T, Messer PW, Petrov DA. 2010. Evidence that Adaptation in Drosophila Is Not Limited by Mutation at Single Sites. PLOS Genetics. 6:1–10.

Kelleher J, Thornton KR, Ashander J, Ralph PL. 2018. Efficient pedigree recording for fast population genetics simulation. PLOS Computational Biology. 14:1–21.

Kim Y, Maruki T. 2011. Hitchhiking Effect of a Beneficial Mutation Spreading in a Subdivided Population. Genetics. 189:213– 226.

Kim Y, Stephan W. 2002. Detecting a Local Signature of Genetic Hitchhiking Along a Recombining Chromosome. Genetics. 160:765–777.

Kimura M. 1957. Some Problems of Stochastic Processes in Genetics. The Annals of Mathematical Statistics. 28:882–901.

Kimura M. 1962. On the probability of fixation of mutant genes in a population. Genetics. 47:713–719.

Kimura M, Ohta T. 1973. The age of a neutral mutant persisting in a finite population. Genetics. 75:199–212.

Kimura M, Weiss GH. 1964. The stepping stone model of population structure and the decrease of genetic correlation with distance. Genetics. 49:561–576.

Lynd A, Weetman D, Barbosa S, Egyir Yawson A, Mitchell S, Pinto J, Hastings I, Donnelly MJ. 2010. Field, Genetic, and Modeling Approaches Show Strong Positive Selection Acting upon an Insecticide Resistance Mutation in Anopheles gambiae s.s. Molecular Biology and Evolution. 27:1117–1125.

Maynard Smith J. 1971. What use is sex? Journal of Theoretical Biology. 30:319–335.

Mazzucco R, Doebeli M, Dieckmann U. 2018. The influence of habitat boundaries on evolutionary branching along environmental gradients. Evolutionary Ecology. 32:563–585.

Messer PW, Neher RA. 2012. Estimating the Strength of Selective Sweeps from Deep Population Diversity Data. Genetics. 191:593–605.

Messer PW, Petrov DA. 2013. Population genomics of rapid adaptation by soft selective sweeps. Trends in Ecology & Evolution. 28:659–669.

Midega JT, Mbogo CM, Mwambi H, Wilson MD, Ojwang G, Mwangangi JM, Nzovu JG, Githure JI, Yan G, Beier JC. 2007. Estimating Dispersal and Survival of Anopheles gambiae and Anopheles funestus Along the Kenyan Coast by Using Mark–Release–Recapture Methods. Journal of Medical Entomology. 44:923–929.

Miles A, pyup.io bot, Murillo R, Ralph P, Kelleher J, Schelker M, Pisupati R, Rae S, Millar T. 2023. scikit-allel 1.3.7. Last accessed 2023-12-28.

Min J, Gupta M, Desai MM, Weissman DB. 2022. Spatial structure alters the site frequency spectrum produced by hitchhiking. Genetics. 222:iyac139.

Muralidhar P, Veller C. 2022. Dominance shifts increase the likelihood of soft selective sweeps. Evolution. 76:966–984.

Nait Saada J, Kalantzis G, Shyr D, Cooper F, Robinson M, Gusev A, Palamara PF. 2020. Identity-by-descent detection across 487,409 british samples reveals fine scale population structure and ultra-rare variant associations. Nature Communications. 11:6130.

Nei M, Li WH. 1979. Mathematical model for studying genetic variation in terms of restriction endonucleases. Proceedings of the National Academy of Sciences. 76:5269–5273.

Nordborg M. 2000. Linkage Disequilibrium, Gene Trees and Selfing: An Ancestral Recombination Graph With Partial Self-Fertilization. Genetics. 154:923–929.

Paulose J, Hermisson J, Hallatschek O. 2019. Spatial soft sweeps: Patterns of adaptation in populations with long-range dispersal. PLOS Genetics. 15:1–29.

Peter BM, Huerta-Sanchez E, Nielsen R. 2012. Distinguishing between Selective Sweeps from Standing Variation and from a De Novo Mutation. PLOS Genetics. 8:1–14.

Prezeworski M, Coop G, Wall JD. 2005. The signature of positive selection on standing genetic variation. Evolution. 59:2312–2323.

Pritchard JK, Pickrell JK, Coop G. 2010. The Genetics of Human Adaptation: Hard Sweeps, Soft Sweeps, and Polygenic Adaptation. Current Biology. 20:R208–R215.

Purcell S, Chang C. 2022. PLINK v1.90b6.26. Last accessed 2023-11-21.

Ralph P, Coop G. 2013. The Geography of Recent Genetic Ancestry across Europe. PLOS Biology. 11:1–20.

Ralph P, Thornton K, Kelleher J. 2020a. Efficiently Summarizing Relationships in Large Samples: A General Duality Between Statistics of Genealogies and Genomes. Genetics. 215:779–797.

Ralph P, Thornton K, Kelleher J. 2020b. tskit 0.5.3. Last accessed 2023-11-21.

Ringbauer H, Coop G, Barton NH. 2017. Inferring Recent Demography from Isolation by Distance of Long Shared Sequence Blocks. Genetics. 205:1335–1351.

Rogers RL, Grizzard SL, Garner JT. 2023. Strong, Recent Selective Sweeps Reshape Genetic Diversity in Freshwater Bivalve Megalonaias nervosa. Molecular Biology and Evolution. 40:msad024.

Roze D. 2016. Background Selection in Partially Selfing Populations. Genetics. 203:937–957.

Sasaki A. 1997. Clumped Distribution by Neighbourhood Competition. Journal of Theoretical Biology. 186:415–430.

Sattath S, Elyashiv E, Kolodny O, Rinott Y, Sella G. 2011. Pervasive Adaptive Protein Evolution Apparent in Diversity Patterns around Amino Acid Substitutions in Drosophila simulans. PLOS Genetics. 7:1–6.

Schrider DR, Kern AD. 2016. S/HIC: Robust Identification of Soft and Hard Sweeps Using Machine Learning. PLOS Genetics. 12:1–31.

Schrider DR, Kern AD. 2017. Soft Sweeps Are the Dominant Mode of Adaptation in the Human Genome. Molecular Biology and Evolution. 34:1863–1877.

Sella G, Petrov DA, Przeworski M, Andolfatto P. 2009. Pervasive Natural Selection in the Drosophila Genome? PLOS Genetics. 5:1–13.

Sianta SA, Peischl S, Moeller DA, Brandvain Y. 2022. The efficacy of selection may increase or decrease with selfing depending upon the recombination environment. Evolution. 77:394–408.

Smith JM, Haigh J. 1974. The hitch-hiking effect of a favourable gene. Genetics Research. 23:23–35.

Städler T, Haubold B, Merino C, Stephan W, Pfaffelhuber P. 2009. The Impact of Sampling Schemes on the Site Frequency Spectrum in Nonequilibrium Subdivided Populations. Genetics. 182:205–216.

Vinatier F, Tixier P, Duyck PF, Lescourret F. 2011. Factors and mechanisms explaining spatial heterogeneity: a review of methods for insect populations. Methods in Ecology and Evolution. 2:11–22.

Wallace B. 1975. Hard and Soft Selection Revisited. Evolution. 29:465–473.

Walsh B, Lynch M. 2018. Evolution and Selection of Quantitative Traits. Oxford University Press.

Wei K, Silva-Arias GA, Tellier A. 2023. Selective sweeps linked to the colonization of novel habitats and climatic changes in a wild tomato species. New Phytologist. 237:1908–1921.

Wright S. 1943. Isolation by distance. Genetics. 28:114–138.

